# Functional Plasticity of the gut and the Malpighian tubules underlies cold acclimation and mitigates cold-induced hyperkalemia in *Drosophila melanogaster*

**DOI:** 10.1101/224832

**Authors:** Gil Y. Yerushalmi, Lidiya Misyura, Heath A. MacMillan, Andrew Donini

## Abstract

At low temperatures, *Drosophila*, like most insects, lose the ability to regulate ion and water balance across the gut epithelia, which can lead to a lethal increase of [K^+^] in the hemolymph (hyperkalemia). Cold-acclimation, the physiological response to low temperature exposure, can mitigate or entirely prevent these ion imbalances, but the physiological mechanisms that facilitate this process are not well understood. Here, we test whether plasticity in the ionoregulatory physiology of the gut and Malpighian tubules of *Drosophila* may aid in preserving ion homeostasis in the cold. Upon adult emergence, *D. melanogaster* females were subjected to seven days at warm (25°C) or cold (10°C) acclimation conditions. The cold acclimated flies had a lower critical thermal minimum (CT_min_), recovered from chill coma more quickly, and better maintained hemolymph K+ balance in the cold. The improvements in chill tolerance coincided with increased Malpighian tubule fluid secretion and better maintenance of K^+^ secretion rates in the cold, as well as reduced rectal K^+^ reabsorption in cold-acclimated flies. To test whether modulation of ion-motive ATPases, the main drivers of epithelial transport in the alimentary canal, mediate these changes, we measured the activities of Na^+^-K^+^-ATPase and V-type H^+^-ATPase at the Malpighian tubules, midgut, and hindgut. Na^+^/K^+^-ATPase and V-type H^+^-ATPase activities were lower in the midgut and the Malpighian tubules of cold-acclimated flies, but unchanged in the hindgut of cold acclimated flies, and were not predictive of the observed alterations in K^+^ transport. Our results suggest that modification of Malpighian tubule and gut ion and water transport likely prevents cold-induced hyperkalemia in cold-acclimated flies and that this process is not directly related to the activities of the main drivers of ion transport in these organs, Na^+^/K^+^- and V-type H^+^-ATPases.

**Summary Statement:** At low temperatures, *insects* lose the ability to regulate ion and water balance and can experience a lethal increase in hemolymph [K^+^]. Previous exposure to low temperatures can mitigate this effect and improve chill tolerance. Here, we show that plasticity of ion and fluid transport across the Malpighian tubule and rectal epithelia likely drive this response.

## 1. Introduction

### 1.1 Chill susceptibility in insects

Chill-susceptible insects are those that succumb to the effects of chilling at temperatures well above the freezing point of their body fluids (Bale, 1996; Overgaard and MacMillan, 2017). While most insect species are considered chill susceptible, the physiology underlying chill susceptibility remains poorly understood. Three metrics are principally used to assess the cold tolerance of chill susceptible insects. The critical thermal minimum (CT_min_) is the temperature at which insects lose coordination and subsequently enter a state of complete neuromuscular paralysis known as a chill coma (Block, 1990; Mellanby, 1939). In the case of mild and/or short cold exposures, chill susceptible insects may recover from a chill coma and regain full neuromuscular function and the time it takes for an insect to stand following removal from a cold exposure is termed chill coma recovery time (CCRT) (Jean David et al., 1998; Macdonald et al., 2004; MacMillan and Sinclair, 2011). Following an intense cold exposure (longer duration and/or lower temperature), chill-susceptible insects acquire irreversible injuries and eventually die (Koštál et al., 2004; Koštál et al., 2006; Rojas and Leopold, 1996) and rates of survival following chilling are also regularly used as a measure of insect cold tolerance (Andersen et al., 2015; MacMillan et al., 2015a; Sinclair et al., 2015). While there is often a high degree of covariance in these three metrics, the mechanisms that underlie each of them is different and uniquely informative, and thus all three are regularly used (Sinclair et al., 2015).

### 1.2 Physiology of chill coma and injury

The resting membrane potential of dipteran muscles and nerves is energized by Na^+^/K^+^-ATPase in two ways: (1) the active transport of cations to the external environment, and (2) the passive leak of positive charge (primarily K^+^) to the extracellular environment, facilitated by the ion gradients set by Na^+^/K^+^-ATPase (Fitzgerald et al., 1996; Thomas, 1972). As such, one of the major challenges for animals at low temperatures is the reduction of enzymatic activity, particularly of ion-motive ATPases, which results in cellular depolarization (Ellory and Willis, 1982; MacMillan et al., 2015b). Cold-induced depolarization thus occurs in two phases, first from the immediate reduction of active ion transport and second from the gradual loss of the ionic gradient necessary for passive ion leak. During the first phase, depolarizations at the onset of a cold exposure have been shown in both muscles and nerves immediately at the onset of a cold exposure (Goller et al., 1990; Hosler et al., 2000; MacMillan et al., 2014). For example, when exposed to low temperatures, various chill-susceptible insects, including *D. melanogaster*, experience muscular depolarization, likely resulting from the reduced electrogenic current of ion-motive ATPases (Goller et al., 1990; Hosler et al., 2000; MacMillan et al., 2014). In addition to reduced electrogenic current, the small extracellular space surrounding locust nerves allows for a rapid surge of extracellular [K^+^] which also likely contributes to initial cellular depolarizations (Robertson et al., 2017; Rodgers et al., 2010). Ultimately reductions of nerve and muscle excitability both likely underlie the neuromuscular paralysis of a chill-coma (Goller et al., 1990; Hazell and Bale, 2011; Hosler et al., 2000).

In contrast to the immediate effects of low temperatures, prolonged cold exposure is often accompanied by large disruptions of hemolymph ion and water homeostasis in many chill-susceptible insects including *Drosophila* (Andersen et al., 2013; Koštál et al., 2004; Koštál et al., 2006; MacMillan and Sinclair, 2011; MacMillan et al., 2015c). When chill susceptible insects are exposed to low temperatures, Na^+^ leaks down its concentration gradient, away from the hemolymph and into the gut (Koštál et al., 2004; MacMillan and Sinclair, 2011). Since Na^+^ is a major hemolymph osmolyte, water passively follows into the gut, leading to an overall reduction in hemolymph water content (Koštál et al., 2004; Koštál et al., 2006; MacMillan et al., 2012). This reduction in hemolymph volume leads to the concentration of hemolymph K^+^, a commonly reported consequence of low temperatures in chill-susceptible insects (Koštál et al., 2004; Koštál et al., 2006; MacMillan et al., 2015c; Yerushalmi et al., 2016). There is also growing evidence suggesting that cold-induced hyperkalemia also stems from the direct leak of K^+^ from tissues and/or the gut in crickets (Des Marteaux and Sinclair, 2016), fruit flies (MacMillan et al., 2015a), and migratory locusts (Andersen et al., 2013; Findsen et al., 2013), but what causes this leak remains unknown. Increased cold-induced hyperkalemia has been linked to longer CCRT and decreased chilling survival, suggesting that failure of ion regulation is a central problem for chill-susceptible insects in the cold (Koštál et al., 2004; Koštál et al., 2006; MacMillan et al., 2015a; Yerushalmi et al., 2016). Chill coma recovery is therefore thought to depend on the restoration of homeostatic hemolymph [K^+^], evidenced by the active reuptake of Na^+^ from the gut into the hemolymph in crickets, which restores hemolymph volume and consequently homeostatic hemolymph [K^+^] (MacMillan et al., 2012). Chilling injury is also closely associated with the degree of hemolymph [K^+^] elevation, such that the time of an approximate two-fold increase in hemolymph [K^+^] is roughly predictive of a species’ median lethal temperature (LT_50_) (Koštál et al., 2004; Koštál et al., 2006; MacMillan and Sinclair, 2011; MacMillan et al., 2014). Cold-induced hyperkalemia further depolarizes resting membrane potential by reducing the K^+^ gradient necessary for passive K^+^ leak, and it is these cumulative short- and long-term depolarizations that has been linked to cellular damage in locusts (MacMillan et al., 2015c).

The chill tolerance of insects (including species of the genus *Drosophila*) can vary as a result of evolutionary adaptation, thermal acclimation, or even acute exposure to low temperatures (rapid cold-hardening) (Andersen et al., 2017b; Chown and Terblanche, 2006; Colinet and Hoffmann, 2012; Hoffmann et al., 2003; Kelty and Lee, 2001; Koštál et al., 2004; Koštál et al., 2006; MacMillan et al., 2015a). For example, exposing *D. melanogaster* to 15°C for six-days as adults extended the Lt_50_ (lethal time of exposure to −2°C that results in 50% survival) from less than 5 h to over 20 h (MacMillan et al., 2015a). These gains in chill tolerance with cold acclimation are closely associated with improved maintenance of ion and water balance in the cold. Together, this body of evidence suggests that failure to maintain water and ion homeostasis in the cold underlies *D. melanogaster* cold tolerance, and that cold acclimation mitigates the extent of this ion imbalance.

### 1.3 Ion and water regulation in insects

At the organismal level, ion and water homeostasis are principally maintained by the transport and permeability of ions and water across the ionoregulatory epithelia, namely the midgut, Malpighian tubules, and hindgut epithelia. It is across these epithelia that cold-induced ion and water leak occurs in crickets (MacMillan and Sinclair, 2011; MacMillan et al., 2012), and thus an investigation of their transport properties before and after thermal acclimation is likely key to understanding cold acclimation.

The midgut is the largest segment of the *Drosophila* gut and is responsible for carrying out the vital functions of nutrient digestion, absorption, and defence against ingested pathogens (Overend et al., 2016). Following the midgut are the Malpighian tubules, diverticula of the gut that function as the main site of ion and water secretion from the hemolymph into the gut in insects. The Malpighian tubules actively transport ions from the hemolymph into the tubule lumen, which osmotically drags water to produce an isosmotic primary urine (Dow and Davies, 2001; Larsen et al., 2014). The primary urine exits the Malpighian tubules where it enters the gut lumen at the junction of the midgut and hindgut mixing with contents from the midgut before passing posteriorly to the hindgut where the reabsorption of water, ions, and metabolites takes place prior to the excretion of wastes (Phillips et al., 1987; Wigglesworth, 1932). The hindgut of *Drosophila* is composed of the ileum and rectum. Most ion and water reabsorption occurs at specialized areas of thickened rectal epithelia called rectal pads that actively absorb ions to create local osmotic gradients for the reabsorption of water (Larsen et al., 2014).

To date, differences in ionoregulatory organ function that relate to cold tolerance have only been described in the Malpighian tubules and rectal pads among *Drosophila* species that differ in chill tolerance (Andersen et al., 2017b; MacMillan et al., 2015d) and in the Malpighian tubules of cold acclimated *G. pennsylvanicus* (Des Marteaux et al., 2018). Whereas low temperatures disturbed the ratio of Na^+^:K^+^ secreted by the tubules of chill susceptible *Drosophila species*, tolerant species experience little or no such change (MacMillan et al., 2015d). Since Na^+^ and K^+^ are the main cations secreted by the tubules, the ratio of Na^+^:K^+^ secretion is particularly informative in illustrating the ion-selective effect of low temperature exposure. Maintenance of this ratio may assist in preventing hyperkalemia by (1) maintaining K^+^ excretion in the cold or (2) minimizing excretion of hemolymph Na^+^, an osmolyte important to the maintenance of hemolymph volume. Further, the rectal pads of chill tolerant *Drosophila* species reabsorb less K^+^ in the cold than those of chill susceptible species, which would also facilitate the maintenance of low hemolymph [K^+^] (Andersen et al., 2017b).

While the underlying mechanisms for the ionoregulatory changes of cold tolerant flies at low temperatures remain unclear, the regulation of ion-motive ATPases that energize these tissues is a possibility. For example, increased ion-motive ATPase activity in the Malpighian tubules would raise the basal fluid and ion secretion rate, allowing greater K^+^ clearance in the cold. Conversely, in absorptive tissues such as the hindgut and the Malpighian tubules, reductions of ion-motive ATPases would reduce ion absorption and thus mitigate the active reuptake of K^+^ in the cold. To date, whole body Na^+^/K^+^-ATPase activity was measured in cold-acclimated Drosophila (MacMillan et al., 2015b) and an organ-specific assessment of Na^+^/K^+^-ATPase and V-type H^+^-ATPase activity in cold-acclimated G. pennsylvanicus_have been conducted (Des Marteaux et al., 2018). In the present study a complete assessment of both organ-specific Na^+^/K^+^-ATPase and V-type H^+^-ATPase activities alongside functional measurements for the midgut, the Malpighian tubules, and the hindgut are presented for *D. melanogaster*.

### 1.4 Experimental goals and hypotheses

In this study, we investigate the effect of cold acclimation following seven days at 10°C on functional ion transport parameters of the ionoregulatory organs of *D. melanogaster*. We hypothesize that both Na^+^/K^+^-ATPase and V-type H^+^-ATPase activities will increase in the Malpighian tubules of cold-acclimated flies, and that these increases will enable a higher capacity for K^+^ clearance at low temperatures (Figure 1). Conversely, we hypothesized that in the midgut and hindgut, decreases in Na^+^/K^+^-ATPase and V-type H^+^-ATPase activities will reduce K^+^ absorption and mitigate hyperkalemia in the cold (Figure 1). To test this, we first confirm that cold-acclimation improves the cold tolerance of *D. melanogaster* and mitigates cold-induced hyperkalemia. We then measure ion transport parameters of Malpighian tubules and gut function directly. Lastly, we quantify the activities of Na^+^/K^+^-ATPase and V-type H^+^-ATPase in the primary ionoregulatory organs (midgut, Malpighian tubules, and hindgut) to test whether their modulation underlies organ-specific function.

**Figure 1.**
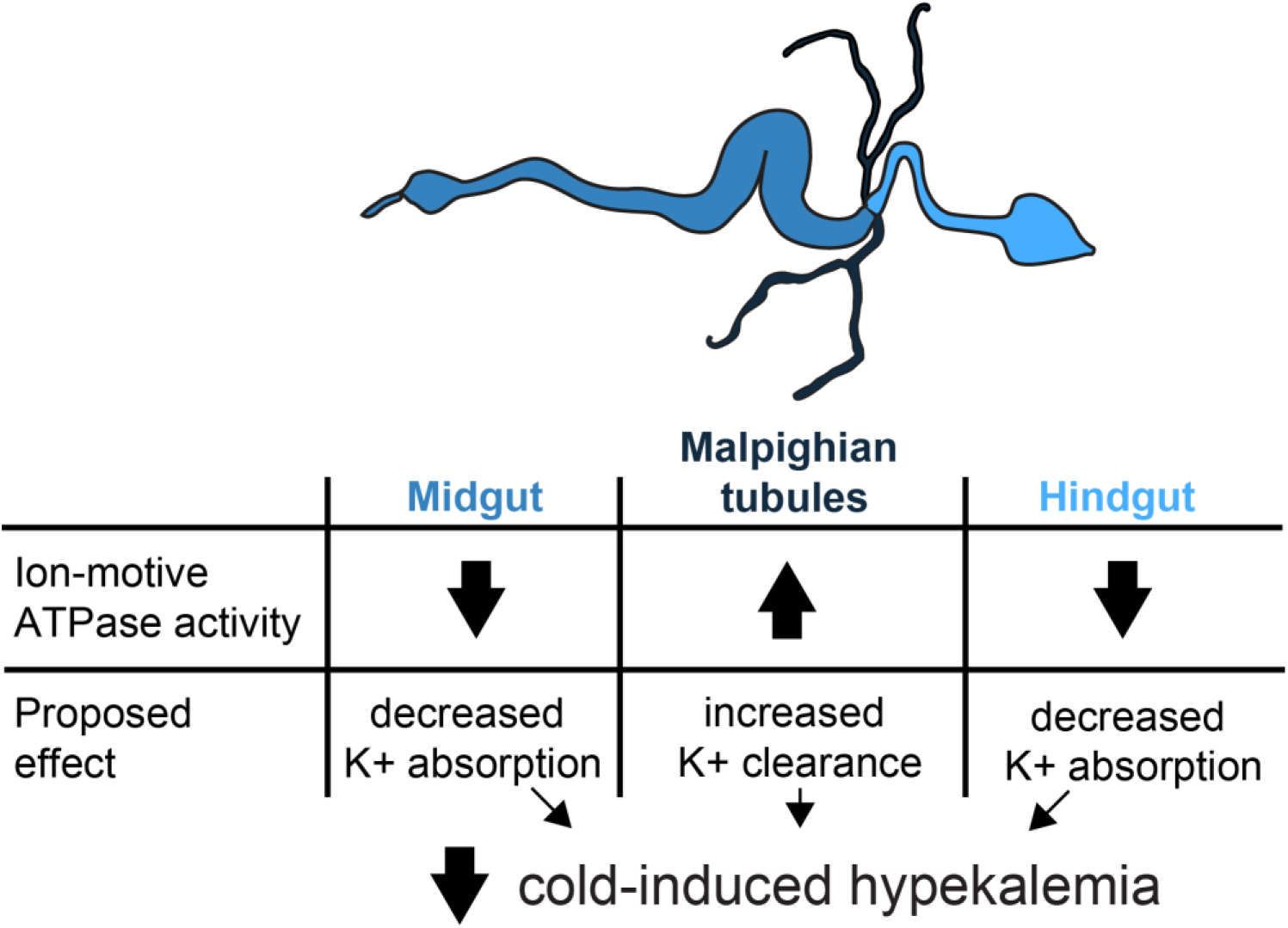
A proposed model of the ionoregulatory changes in the midgut, Malpighian tubules, and hindgut that prevent cold-induced hyperkalemia in cold-acclimated *D. melanogaster*. Ion-motive ATPases such at Na^+^/K^+^-ATPase and V-type H^+^-ATPase are the main drivers of epithelial transport in the gut and the Malpighian tubules of insects, and their activity is proposed to alter gut and tubule function to reduce K^+^ absorption and increase K^+^ excretion. Decreased ion-motive ATPase activity in absorptive organs such as the midgut and the hindgut is therefore predicted to reduce K^+^ absorption while increased ion-motive ATPase activity in the Malpighian tubules is proposed to increase K^+^ clearance. Cumulatively, these changes are proposed to facilitate net K^+^ excretion and the maintenance of low hemolymph [K^+^] in the cold.

## 2. Methods

### 2.1 Animal husbandry and acclimation treatments

The population of *Drosophila melanogaster* used in this study was established in 2008 by combining 35 isofemale lines from southwestern Ontario, Canada (Marshall and Sinclair, 2010). Fly rearing was conducted as previously described (Yerushalmi et al., 2016) by transferring mature adults into 200 mL plastic bottles containing ~50 mL of a standard rearing diet (Bloomington *Drosophila* medium; Lakovaara, 1969) for 1-2 h, ensuring an approximate egg density of 100-150 eggs/bottle. The bottles were then stored at 25°C and a 14:10 h light:dark cycle. Filter paper was placed in each bottle to increase surface area for pupation. Newly-emerged adults were collected daily and transferred into 40 mL plastic vials containing 7-10 mL of the rearing diet. The vials were then randomly assigned to one of the two treatments: warm- or cold-acclimation. Warm-acclimated (WA) flies were maintained at 25°C with a 14:10 h light:dark cycle and cold-acclimated (CA) flies were maintained at 10°C with 10:14 h light:dark cycle (aimed to mimic summer and fall conditions, respectively). All experiments were conducted on non-virgin adult females following a one-week exposure to their acclimation treatment.

### 2.2 Chilling tolerance phenotypes

In the present study we measured the critical thermal minimum (CT_min_) and chill coma recovery time (CCRT). Chilling survival was recently measured in the same laboratory population of flies under identical rearing and acclimation conditions, and was described by MacMillan et al. (2017).

To measure CT_min_ as previously described (Andersen et al., 2015; Yerushalmi et al., 2016), flies were individually placed in 4 mL glass screw-top vials. Vials were then attached to a custom-built rack and placed in a temperature-controlled bath (Model MX7LL, VWR International, Mississauga, Canada) containing a 1:1 mixture of ethylene glycol and water at room temperature (25°C). The bath temperature was then ramped down at a rate of −0.15°C min^−1^ and the temperature was monitored independently using a pair of type-K thermocouples connected to a computer running Picolog (version 5.24.8) via a Pico TC-08 interface (Pico Technology, St. Neots, UK). Flies were individually observed throughout the ramping period and the temperature at which no fly movement was observed following a disturbance of the vial with a plastic probe was recorded as its CT_min_.

To measure CCRT as previously described (MacMillan et al., 2015a; Yerushalmi et al., 2016), female flies were individually placed in 4 mL glass screw-top vials. The vials were sealed in a plastic bag and submerged in an ice-water mixture (0°C). After 6h the vials were removed from the ice-water mixture and placed at room temperature (25°C) where the flies were individually observed. The time that it took an individual fly to stand on all six legs following its removal from the cold treatment was recorded as its CCRT.

### 2.3 Hemolymph [K^+^] measurements

Hemolymph [K^+^] was assessed in flies from both acclimation groups following exposure to 0°C for various durations from 0 h to the maximal survival duration at 0°C (up to 30 h for warm-acclimated flies and up to 110 h for cold acclimated flies) using the ion-selective microelectrode technique (ISME) as previously described (Jonusaite et al., 2011). The cold-exposure durations used here mirrored those previously used in a chill-survival analysis (MacMillan et al., 2017), except for 36 h and 42 h for warm-acclimated flies, where hemolymph extraction was unsuccessful and survival rates were very low. Hemolymph droplets were collected by placing individual flies in 200 μL pipette tips and attaching the tips to a custom-made device (MacMillan and Hughson, 2014). Air pressure was applied to position the fly head at the end of a pipette tip and an antenna was then carefully removed under a dissection microscope. Droplets of hemolymph that emerged at ablated antenna were immediately placed under paraffin oil for assessment using ISME.

Ion-selective microelectrodes were prepared from borosilicate glass capillaries (TW150-4; World Precision Instruments, Sarasota, USA) and pulled using the P-97 Flaming-Brown micropipette puller (Sutter Instruments Co., Novato, USA) to a tip diameter of ~5 μm. The microelectrodes were then salinized in vapours of N,N-dimethyltrimethylsilylamine (Fluka, Buchs, Switzerland) at 300°C for 1 hour. K^+^-selective microelectrodes were backfilled with 100 mM KCl and front loaded with the K^+^ Ionophore cocktail B (100 mM KCl; Fluka). Na^+^-selective electrodes (used for assessment of [Na^+^] in the Malpighian tubule secreted fluid, see below) were backfilled with 100 mM NaCl and front loaded with the Na^+^ Ionophore II cocktail A (100 mM KCl/100 mM sodium citrate, pH 6.0; Fluka). The ion-selective microelectrodes were then dipped in polyvinyl chloride (PVC) to prevent the leakage of ionophore into the paraffin oil. To complete the circuit, a conventional microelectrode was prepared from borosilicate glass capillaries (IB200F-2; WPI) and backfilled with 500 mM KCl. Both electrodes were connected to the PowerLab 4/30 data acquisition system (AD Instruments Inc., Colorado Springs, USA) through an ML 165 pH Amp and analyzed with LabChart 6 Pro software (AD Instruments Inc.). Once the set up of ISME was complete, 5 μL calibration droplets with known concentrations of the ion of interest and a 10-fold difference in its concentration were measured. To ensure ion-specificity, the lower of the two concentration droplets was corrected in ionic strength using LiCl. For example, for the measurement of hemolymph [K^+^], 10 mM KCl/90 mM LiCl and 100 mM KCl were used for calibration. The final ion concentrations were then calculated with the following equation:

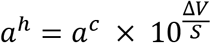

Where a^h^ is the hemolymph ion concentration, a^c^ is the concentration of one of the calibration droplets, ΔV is the difference in voltage between the hemolymph and the calibration solution and S is difference in voltage between two calibration droplets with a tenfold difference in ion activity.

### 2.4 Malpighian tubule fluid and ion secretion rates

To assess differences in Malpighian tubule activity, modified Ramsay assays (Ramsay, 1954) were conducted on tubules extracted from cold- and warm-acclimated flies and at 0°C, 5°C, 10°C, and 23°C. To isolate Malpighian tubules, individual flies were first dipped in 70% ethanol for 5-10 seconds to remove the cuticular waxes and transferred into a dish lined with a silicone elastomer (Sylgard 184; Dow Corning Corp., Midland, USA) and containing *Drosophila* saline (10 mM glutamine, 20 mM glucose, 15 mM MOPS, 4.2 mM NaH_2_PO_4_, 10.2 mM NaHCO_3_, 8.5 mM MgCl_2_ (hexahydrate), 2 mM CaCl_2_ (dihydrate), 20 mM KCl, 117.5 mM NaCl, pH 7.0) for dissections. The anterior pair of Malpighian tubules were then isolated along with a portion of the ureter by cutting the ureter near the ureter-gut junction. Upon their removal, the Malpighian tubules were transferred into another silicone-lined dish containing 35 μL droplets of a 1:1 mixture of *Drosophila* saline and Schneider’s insect medium (Sigma-Aldrich, St. Louis, USA) placed in premade wells under paraffin oil. Using a glass probe, one tubule was placed in the droplet and the other was carefully wrapped around a metal pin adjacent to the droplet, ensuring exposure of the excised ureter to the paraffin oil where secreted fluid would accumulate to be collected for analysis (see Larsen et al., 2014).

To assess Malpighian tubule function at 0°C, 5°C and 10°C, after the Ramsay assays were set up at room temperature, the silicone-lined dish housing the assay was placed in a glass dish containing ~1cm of water within a Precision™ Low Temperature BOD Refrigerated Incubator (Model PR205745R, ThermoFisher Scientific, Waltham, USA). The temperature of the water bath holding the Ramsay assay dishes was monitored independently using type-K thermocouples and maintained within 1°C of treatment temperature at all times. Because chilling slows rates of fluid transport (Anstee et al., 1979; MacMillan et al., 2015d; Maddrell, 1964), incubation times were adjusted depending on temperature to ensure a droplet of sufficient size for measurement and analysis by ISME; Ramsay assays were incubated for 30 min, 120 min, 150 min, or 180 min for assays running at 23°C, 10°C, or 5°C, or 0°C, respectively. Following this incubation period, a glass probe was used to isolate the primary urine droplet under paraffin oil. Droplets of the secreted fluid were then suspended in oil (to ensure a spherical shape) and droplet diameter was measured using the ocular micrometer of a Motic^®^ K-400L Stereo Microscope (Motic North America, Richmond, Canada). The fluid secretion rate was then calculated using the following equation:

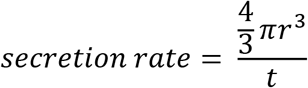

Where r is the radius of the secreted fluid droplet in mm, and t is the incubation period of the Ramsay assay in minutes.

Lastly, [Na^+^] and [K^+^] of the fluid secreted by the Malpighian tubules were measured using ISME as described above (see section 3.3).

### 2.5 Midgut and hindgut K^+^ flux

The scanning ion-selective microelectrode technique (SIET) was used to measure K^+^ flux across the midgut and hindgut epithelium as previously described (Andersen et al., 2017c; Jonusaite et al., 2013; Rheault and O’Donnell, 2001). Briefly, a K^+^-selective microelectrode was prepared as described above and mounted onto a headstage with an Ag/AgCl wire electrode (WPI). The headstage was connected to an ion polarographic amplifier (IPA-2, Applicable Electronics, Forestdale, USA). The circuit was completed using a reference electrode composed of 3% agar in 3 M KCl that solidified inside a glass microcapillary. One end of the electrode was placed in the bathing solution while the other end was connected to a headstage via an Ag/AgCl half-cell (WPI). Ion selective microelectrodes were calibrated in 5 mM KCl/45 mM LiCl and 50 mM KCl solutions.

Rates of K^+^ flux at the midgut and rectum of both warm- and cold-acclimated flies were measured at 23°C and 6°C using SIET. Whole guts were carefully isolated and bathed in fresh *Drosophila* saline in the lid of 35 mm Petri dishes. The use of the Petri dish lid minimized gut movement due to the adhesion of the gut to the surface of the dish. Individual measurements were conducted 5-10 μm from the gut epithelium and 100 μm away, for the assessment of concentration differences near and away from the preparation. To minimize potential gradient disturbance effect of the electrode movement, a 4 s wait time was employed between the two positions of measurement, followed by a 1s recording period. For each measured position, this protocol was repeated four times, and the average voltage gradient between the two points was used for the calculation of K^+^ flux. The Automated Scanning Electrode Technique (ASET) software (version 2.0, Science Wares, East Falmouth, USA) was used to automatically run the sampling protocol and calculate the average voltage gradient at each assessed site. A measurement of background noise was recorded for each preparation ~3mm away from the gut and was used in the calculation of K^+^ flux to account for mechanical disturbances in the ion gradients that arise from the movement of the electrode during sampling.

Voltage gradients were converted into concentration gradients using the following equation as previously described by Donini and O’Donnell (2005):

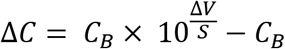

Where ΔC is the concentration gradient between the two measured points, C_B_ is the background gradient measured away from the gut preparation, ΔV is the voltage gradient adjacent to the tissue, and S is the difference in voltage between two calibration droplets with a tenfold difference in ion activity. While in reality this technique measures ion activity, data can be expressed in concentrations if it is assumed that the ion activity coefficient in the experimental solution is the same as that of the calibration (Donini and O’Donnell, 2005).

Measurement sites across the gut included six equidistant sites across the midgut (averaged and represented as a single flux), three sites across the ileum (averaged and represented as a single flux), and 2-3 sites on the rectal pads (averaged and represented as a single flux). For each site, two or more measurements were taken and averaged.

### 2.6 Na^+^/K^+^-ATPase and V-type H^+^-ATPase enzyme activity

Tissue-specific Na^+^/K^+^-ATPase and V-type H^+^-ATPase activities were measured as described by Jonusaite et al. (2011) by quantifying the oubain- (Sigma-Aldrich Canada, Oakville, Canada) or bafilomycin-sensitive (LC Laboratories, Woburn, USA) hydrolysis of adenosine triphosphate (ATP) at 25°C.

Midguts, Malpighian tubules, and hindguts were each collected from warm- and cold-acclimated flies. To isolate these organs, individual flies were first dipped in 70% ethanol for 5-10 seconds for the removal of cuticular waxes and transferred into a dish lined with a silicone elastomer containing *Drosophila* saline for dissections. To minimize rapid thermal plasticity, dissections were conducted at temperatures approximating the acclimation treatments. Warm-acclimated flies were dissected at room temperature (~23°C) and cold-acclimated flies were dissected at ~10°C by placing the dissecting dish on a PE100 Inverted Peltier System connected to a PE95 controller (Linkam Scientific Instruments, Tadworth, England) in the view of the dissecting microscope. Following the dissection of each individual fly, isolated organs were transferred to 2 mL microcentrifuge tubes and immediately flash frozen using liquid nitrogen. Frozen samples were stored at −80°C for later tissue processing.

To homogenize the organs, samples were thawed on ice and 100 μL of homogenizing buffer was added to each tube (150 mM sucrose,10 mM Na_2_EDTA, 50 mM imidazole, 0.1% deoxycholic acid; pH 7.3). The samples were homogenized on ice using a PRO 250 homogenizer with a 5 × 75 mm generator (PRO Scientific Inc., Oxford, USA) for 8-10 seconds and centrifuged at 10,000 × *g* for 10 minutes at 4°C using a 5810R centrifuge (Eppendorf Canada, Mississauga, Canada). The resulting supernatants were then collected into 2 mL tubes and stored at −80°C.

Three assay solutions (A, B, and C) containing the appropriate enzymes and reagents were prepared. Solution A was composed of 4 U/mL lactate dehydrogenase (LDH), 5 U/mL pyruvate kinase (PK), 2.8 mM phosphoenolpyruvate (PEP), 3.5 mM ATP, 0.22 mM NADH, 50 mM imidazole, and a pH of 7.5. Solutions B and C were similar in composition but also contained 5 mM oubain or 10 μM bafilomycin, respectively, for the inhibition of the ATPases under investigation. Following their preparation, each solution was mixed in a 3:1 ratio with a salt solution composed of 189 mM NaCl, 10.5 mM MgCl_2_, 42 mM KCl, 50 mM imidazole, and a pH of 7.5. Final conditions for the assays were as follows: 3 U/mL LDH, 3.75 U/mL PK, 2.1 mM PEP, 2.63 mM ATP, 0.17 mM NADH, 47.25 mM NaCl, 2.6 mM MgCl_2_, 10.5 mM KCl, 50 mM imidazole, and a pH of 7.5

Prior to running the assays, an adenosine diphosphate (ADP) standard curve was run to ensure that all reagents used are working normally. First, 0 nM, 5 nM, 10 nM, 20 nM, 40 nM ADP standards were prepared by diluting stock ADP in imidazole buffer. Two technical replicates containing 10 μL of each ADP standard were then added to a 96-well polystyrene microplate (BD Falcon™, Franklin Lakes, USA) and 200 μL of the assay solution (solution A + salt solution) was added to each well. The plate was then placed in a Thermo Electron Multiskan™ Spectrum microplate spectrophotometer (Thermo Electron Co., Waltham, USA) set to 25°C and measuring absorbance at 340 nm (the peak absorbance of NADH). The recorded absorbance was analyzed using Skanlt version 2.2.237 (Thermo Electron Co.). The assay solution was approved if the optical density of the ADP standards were within 0.2 and 0.9 and if the slope of the curve was between −0.012 and −0.014 OD nmol ADP^−1^ (Jonusaite et al., 2011).

To run both assays (Na^+^/K^+^-ATPase and V-type H^+^-ATPase), experimental homogenates were thawed and added to a 96-well microplate that was kept on ice in six replicates of 10 μL each. Following this, 200 μL of the assay solutions (Salt solution mixed with solutions A, B, or C) were added to two replicates for each experimental sample, resulting in two technical replicates per sample. The plate was then inserted into the microplate reader and the linear disappearance of NADH (peak absorbance: 340 nm) was assessed over a 30 min period.

Upon assay completion, raw absorbance data was extracted from the Multiskan Spectrum data acquisition system and the rates of NADH disappearance were independently assessed in R version 3.3.1 (R Core Team, 2015) using the lmList() function available in the lme4 package (Bates et al., 2016). Na^+^/K^+^-ATPase and V-type H^+^-ATPase specific ATP consumption was determined by assessing the difference in activity between samples running with or without oubain or bafilomycin, respectively. Enzyme activity was standardized to the protein content of each sample using a Bradford assay (Sigma-Aldrich Canada) according to the manufacturer’s guidelines and using bovine serum albumin as a standard (Bioshop Canada Inc., Burlington, Canada). Final enzyme activities were calculated using the following equation:

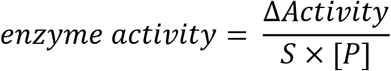

Where Δ*Activity* is the difference in the rate of ATP hydrolysis in the absence and presence of ouabain or bafilomycin, S is the slope of the ADP standard curve, and [P] is the protein concentration of the sample.

### 2.7 Malpighian tubule size

In analysis of ion-motive ATPase activities in the Malpighian tubules, it became apparent that the protein content of cold-acclimated Malpighian tubules was significantly elevated (Figure 6A). Thus, to investigate the cause of this difference in protein content, Malpighian tubule size was assessed for flies of both acclimation groups. The anterior pair of tubules of individual flies were dissected out, ensuring that no direct contact was made with the Malpighian tubules themselves. Images of the tubules were then captured using an Olympus IX81 inverted microscope (Olympus Canada, Richmond Hill, Canada). Images were recorded and analyzed using Olympus cellSens digital imaging software version 1.12 (Olympus Canada). Malpighian tubule length was measured from the ureter-Malpighian tubule junction to the distal end of each tubule. Tubule width measurements were always taken ~100 μm from the ureter-tubule junction.

### 2.8 Statistical analysis

The CT_min_ and CCRT of warm- and cold-acclimated flies were compared using unpaired student’s t-tests. Two-way ANOVAs were used to determine the independent and interacting effects of acclimation temperature and exposure temperature on Malpighian tubule fluid and ion secretion rates, ion concentrations in the secreted fluid, and the ratio of Na^+^:K^+^ in the secreted fluid. Holm-Sidak post-hoc tests were then conducted to compare differences in activity between the two acclimation treatments at each exposure temperature. The effects of exposure temperature and acclimation temperatures on K^+^ flux across the midgut, ileum, and rectum were also analyzed using two-way ANOVAs. An ANCOVA was used to assess the effect of exposure duration on hemolymph [K^+^] and to compare this effect between warm- and cold-acclimated flies. Tissue-specific ATPase activity, Malpighian tubule protein content, and Malpighian tubule length and width were compared between the two acclimation treatments using unpaired student’s t-tests. All statistical tests were conducted on GraphPad Prism version 6.0.1 (GraphPad Software, La Jolla, USA).

## 3. Results

### 3.1 Cold acclimation improved chill tolerance and mitigated cold-induced hyperkalemia

The CT_min_ of cold-acclimated flies was significantly lower than that of warm-acclimated flies (unpaired t-test; *P* < 0.0001; Figure 2A); on average, cold-acclimated individuals entered a chill coma ~3.5°C below warm-acclimated flies. The chill coma recovery time following 6 h at 0°C was lower by approximately 50% in cold-acclimated flies (unpaired t-test; *P* < 0.0001; Figure 2B). While cold-acclimated flies recovered in 17.3 ± 0.9 min, warm-acclimated flies required 36.4 ± 2.9 min to recover from the same amount of time at 0°C.

**Figure 2.**
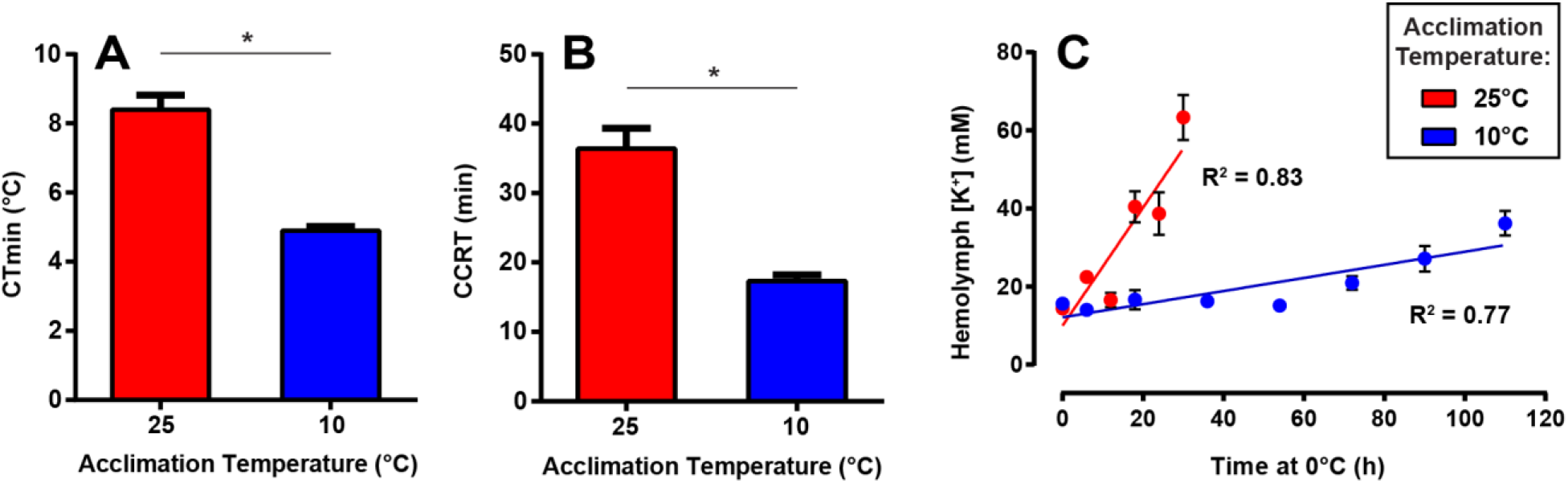
Cold acclimation mitigates cold-induced hyperkalemia and improves the chill tolerance of adult *D. melanogaster* females. (A) Critical thermal minimum (CT_min_) of warm- and cold acclimated flies (n = 18 flies per group). (B) Chill coma recovery time (CCRT) for warm- and cold-acclimated flies (n = 20 flies per group). Both CT_min_ and CCRT were significantly lower in cold-acclimated flies. (C) Hemolymph [K^+^] of cold- and warm-acclimated flies following exposure to 0°C. All bars represent mean ± SEM. Asterisks denote significant difference (unpaired t-test; *P* < 0.001).

Hemolymph [K^+^] levels were assessed in warm- and cold-acclimated flies following exposures to 0°C of varying durations reflecting our previous assessment of chill survival in this population (MacMillan et al., 2017). With increasing duration to cold exposure, hemolymph [K^+^] significantly increased in both warm-acclimated flies (*P* = 0.0045, R^2^ = 0.83) and cold-acclimated flies (*P* = 0.0117; R^2^ = 0.77; Figure 2C). The rate of [K^+^] accumulation, however, differed by a factor of nine among the acclimation groups (ANCOVA; F_1,10_ = 27.27, *P* =

0.0004); whereas hemolymph [K^+^] increased at a rate of ~1.5 mM/hour in warm-acclimated flies, it increased at a rate of ~0.17 mM/hour in cold-acclimated flies (Figure 2C).

### 3.2 Cold acclimation altered Malpighian tubule fluid and ion secretion across a range of temperatures

Malpighian tubule function was measured using Ramsay assays at 0°C, 5°C, 10°C, and 23°C (n = 7-16 individuals per temperature per acclimation group). Exposure temperature and acclimation temperatures interacted to affect all assessments of Malpighian tubule function including fluid secretion rates, ion concentrations in the secreted fluid, ion secretion rates, and the ratio of Na^+^:K^+^ secretion (*P* < 0.05 in all cases; see Table S1 for all two-way ANOVA results). Notably, the fluid secretion rates at 5°C, 10°C, and 23°C were significantly higher in cold-acclimated flies (Holm-Sidak test; *P* = 0.0015, *P* = 0.0002, *P* < 0.0001, respectively; Figure 3A) where the most pronounced difference was an 9-fold higher fluid secretion rate at 10°C, the same temperature as the cold-acclimation temperature (Figure 3A). Cold acclimation altered both [Na^+^] and [K^+^] in the secreted fluid. For warm-acclimated flies, [Na^+^] was relatively stable in the secreted fluid between 5°C and 23°C (~75 mM), but was elevated to 147 ± 19 mM at 0°C (Figure 3B). Cold-acclimated flies secreted fluid with lower [Na^+^] relative to warm-acclimated flies at every tested temperature (Holm-Sidak test; *P* = 0.0032, *P* = 0.0442, *P* = 0.0002, *P* = 0.0128 for 0°C, 5°C, 10°C, and 23°C; Figure 3B), and never exceeded the [Na^+^] measured at 23°C. In parallel, while [K^+^] was stable in cold-acclimated flies throughout all exposure temperatures, [K^+^] in the secreted droplets of warm-acclimated flies was significantly reduced at 0°C and 10°C in comparison to cold-acclimated flies (Holm-Sidak test; *P* = 0.0004, and *P* = 0.0073, respectively; Figure 3C). Changes in Na^+^ and K^+^ concentrations of the fluid secreted by the Malpighian tubules can result from changes to fluid or ion secretion rates, or both. Rates of Na^+^ secretion at 23°C were ~50% higher in cold-acclimated flies, a significant difference (Holm-Sidak test; *P* = 0.0030; Figure 3D). Similar trends were noted at 5°C and 10°C where Na^+^ secretion was increased by 86% and 115%, respectively, but these differences were not significant. While K^+^ secretion decreased with decreasing temperatures in both acclimation groups, cold-acclimated flies maintained higher K^+^ secretion at all temperatures (Holm-Sidak test; *P* = 0.0060, *P* = 0.0018, *P* = 0.0002, *P* = 0.0022, for 0°C, 5°C, 10°C, and 23°C, respectively; Figure 3E). Taken together, these changes in ion transport rates prevented the rise of Na^+^:K^+^ at low temperatures in cold-acclimated flies, where the ratio of Na^+^:K^+^ in the secreted fluid was maintained between 0.18 ± 0.10 at 10°C and 0.56 ± 0.10 at 23°C (Figure 3F). In contrast, this ratio was highly disturbed in warm-acclimated flies, rising from 0.59 ± 0.16 at 23°C to 1.87 ± 0.43 at 0°C. This ratio of Na^+^:K^+^ was significantly higher in warm-acclimated flies in comparison to cold-acclimated flies at 0°C and 10°C (Holm-Sidak test; *P* = 0.0080, *P* = 0.0093, respectively; Figure 3F) but not at 5°C or 23°C.

**Figure 3.**
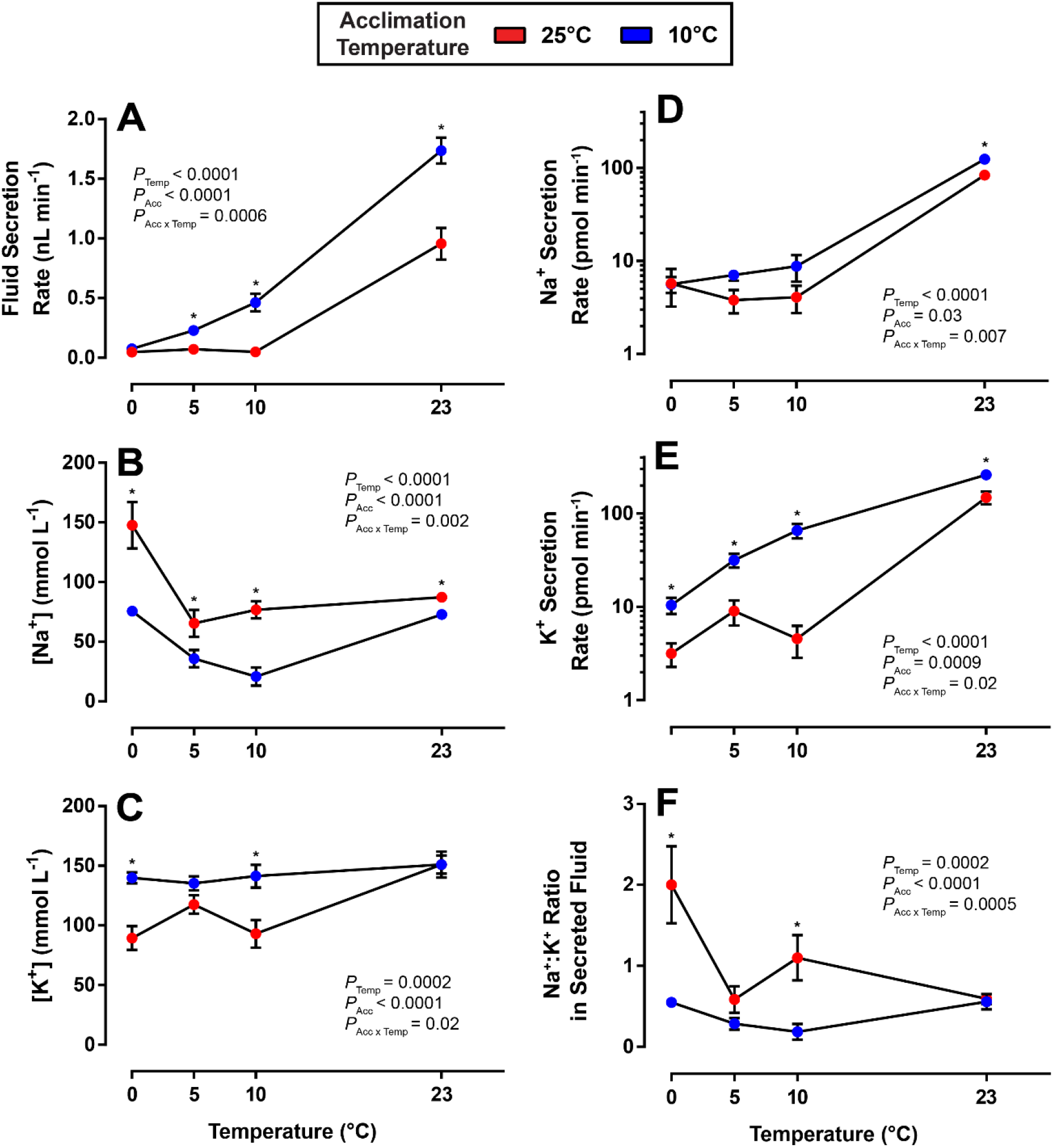
Cold acclimation altered Malpighian tubule function in female *D. melanogaster* across a variety of thermal conditions. (A) Malpighian tubule fluid secretion rate, (B) [K^+^] and (C) Na^+^ in the secreted fluid, (D) Na^+^ and (E) K^+^ secretion rates, and (F) Na^+^:K^+^ secretion ratio assessed at 0°C, 5°C, 10°C, and 23°C in warm- (red) and cold-acclimated flies (blue). Bars represent mean ± SEM, and bars that are not clearly visible are obscured by the symbols. Asterisks denote significant differences between warm- and cold acclimation at the same exposure temperature (Holm-Sidak test; *P* < 0.05). Two-way ANOVAs were conducted on all three variables and the resulting *P*-values are embedded in each respective panel (see Table S1 for all two-way ANOVA results).

### 3.3 Cold-acclimation reduced rectal reabsorption of K^+^

To assess the impact of thermal acclimation on gut K^+^ flux in the cold, K^+^ flux was assessed along the midgut and hindgut (ileum and rectum) at 6°C and 23°C using the scanning ion-selective electrode technique (Figure 4). In the midgut, neither exposure temperature (F_1,20_ = 0.19, *P* = 0.67; Figure 4A) nor acclimation temperature (F_1,20_ = 0.57, *P* = 0.57) altered mean K^+^ flux. In contrast, mean ileal K^+^ flux was predicted by acclimation temperature (F_1,36_ = 4.31, *P* = 0.045; Figure 4B) and its interaction with exposure temperature (F_1,36_ = 6.37, *P* = 0.02), but not exposure temperature (F_1,36_ = 1.30, *P* = 0.26). Lastly, both exposure temperature (F_1,32_ = 4.29, *P* = 0.047) and acclimation temperature (F_1,32_ = 4.33, *P* = 0.046) predicted K^+^ flux at the rectum such that K^+^ flux was lower in cold-acclimated flies and at lower exposure temperatures (Figure 4C).

**Figure 4.**
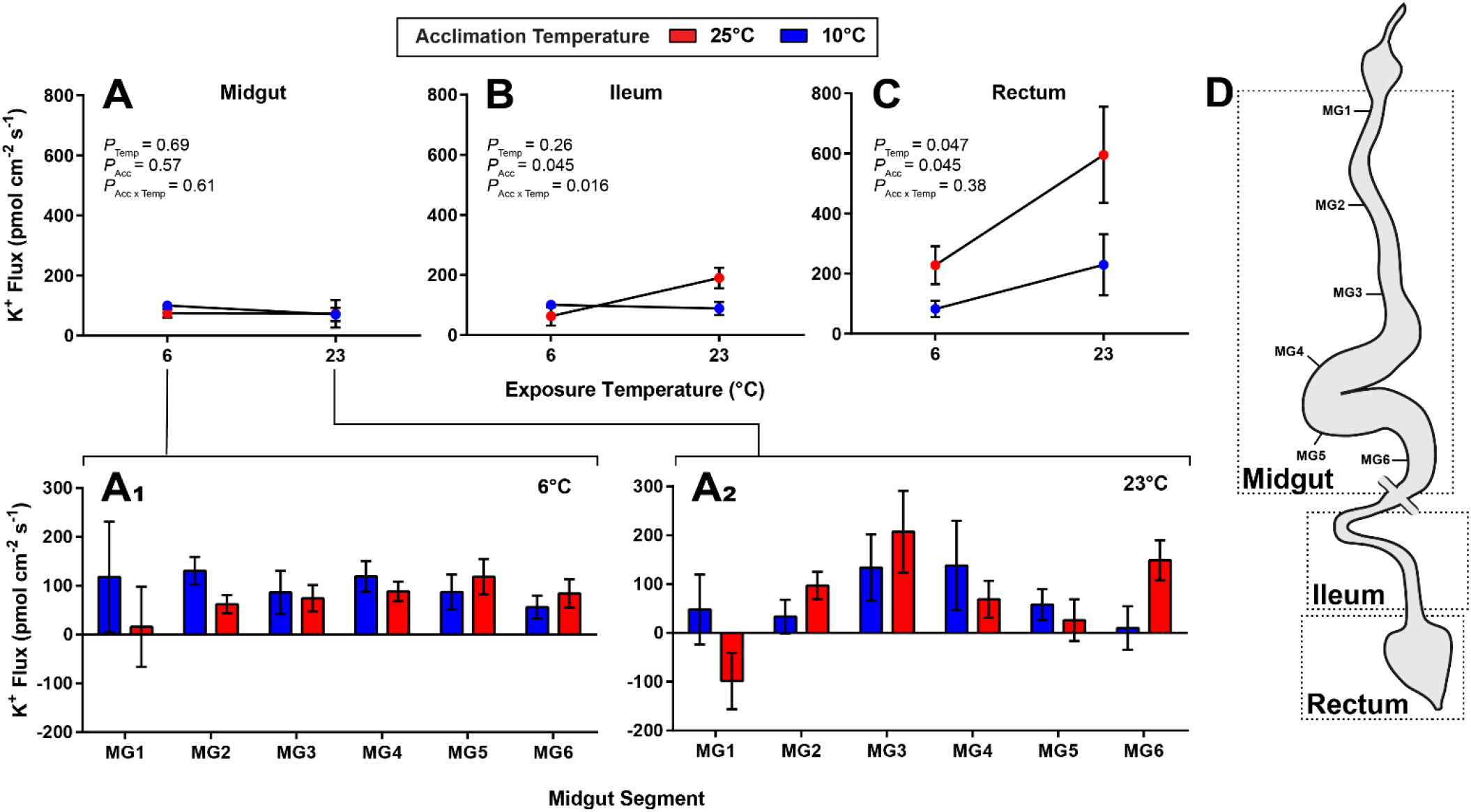
K^+^ reabsorption is reduced in the rectum of cold-acclimated *D. melanogaster* females. (A) Mean K^+^ flux in the midgut (average of six midgut sites), (B) ileum, and (C) rectum (rectal pads) of warm- (red) and cold-acclimated flies (blue). (D) Schematic of alimentary canal illustrating sites of K^+^ flux measurements. Midgut K^+^ flux was measured at six equidistant sites along the midgut denoted MG1 (anterior end) to MG6 (posterior end) at (A_1_) 6°C and (A_2_) 23°C. Bars represent mean ± SEM.

### 3.4 Cold acclimation decreased midgut and Malpighian tubule Na^+^/K^+^-ATPase and V-type H^+^-ATPase activity

The enzymatic activity of Na^+^/K^+^-ATPase and V-type H^+^-ATPase (relative to total protein) was assessed in the midgut, Malpighian tubules, and hindgut of warm- and cold-acclimated flies (Figure 5). Reductions in the maximal activity of both ATPases were noted in the Malpighian tubules and the midgut. Activity of Na^+^/K^+^-ATPase was 41% lower in the midgut (unpaired t-test; *P* = 0.01; n = 5) and 53% lower in the Malpighian tubules (unpaired t-test; *P* = 0.006; n = 3) of cold-acclimated flies relative to those that were warm-acclimated. By contrast, no difference in Na^+^/K^+^-ATPase activity was found in the hindgut (unpaired t-test; *P* = 0.7 n = 5; Figure 5). In a similar pattern, V-type H^+^-ATPase activity was 92% lower in the midgut (unpaired t-test; *P* = 0.0004; n = 3), 61% lower in the Malpighian tubules (unpaired t-test; *P* = 0.01; n = 5) and did not differ in the hindgut of cold-acclimated flies (unpaired t-test; *P* = 0.9; n = 5; Figure 5).

**Figure 5.**
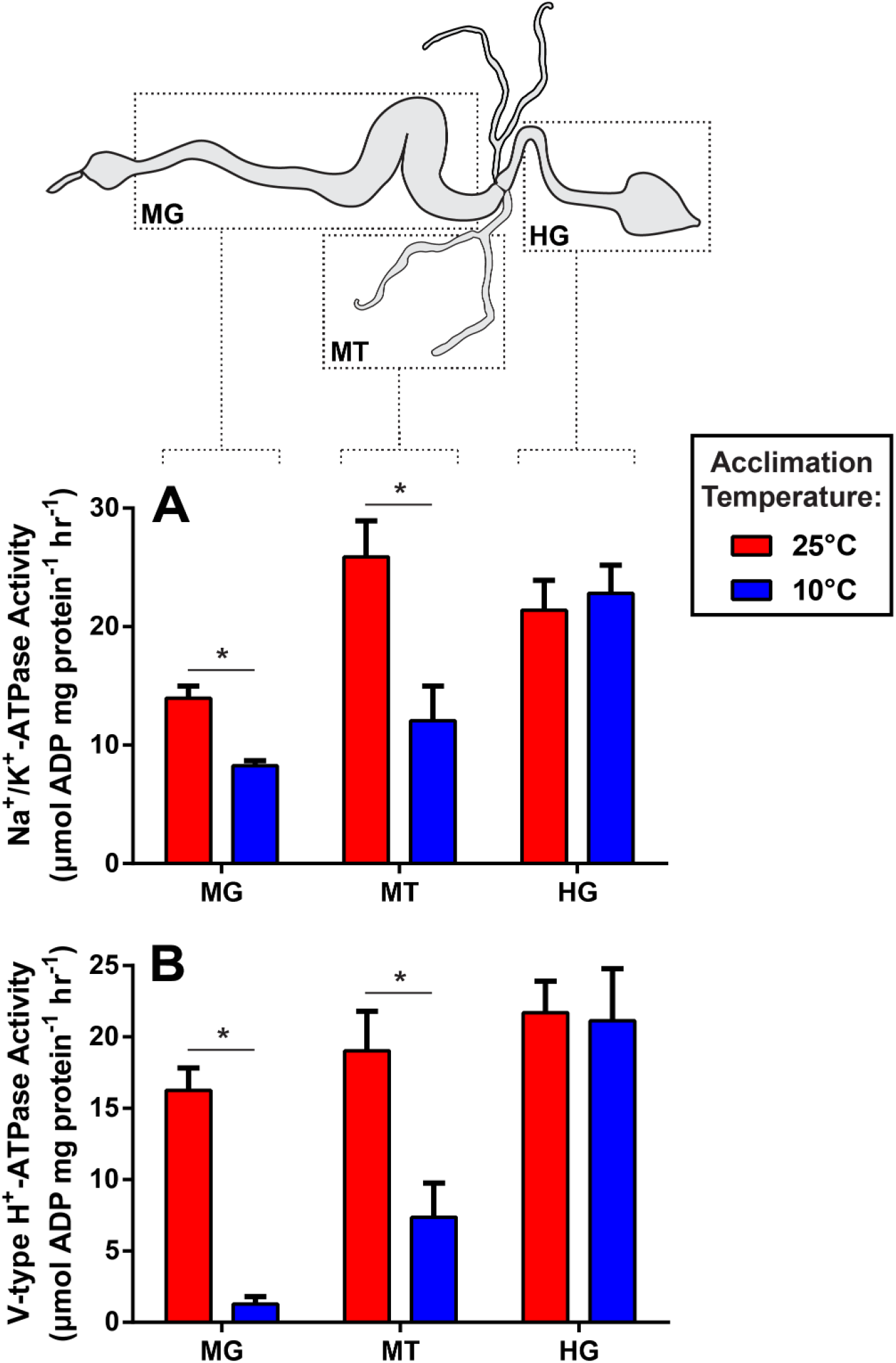
Cold-acclimation decreased the activity of Na^+^/K^+^-ATPase and V-type H^+^-ATPase (relative to protein content) in the midgut and Malpighian tubules of adult female *D. melanogaster*. (A) Enzymatic activity of Na^+^/K^+^-ATPase in the midgut (MG), Malpighian tubules (MT), and hindgut (HG) of warm- (red bars) and cold-acclimated (blue bars) flies. (B) Enzymatic activity of V-type H^+^-ATPase in the midgut, Malpighian tubules, and hindgut in warm- and cold-acclimated flies. Bars represent mean ± SEM. Asterisks denote significant difference in enzymatic activity (unpaired t-tests; *P* < 0.05).

Interestingly, differences in activity of both ATPases in the Malpighian tubules were largely driven by differences in protein content (unpaired t-test; *P* = 0.01; Figure 6A), despite a similar number of Malpighian tubules collected per group. When assessed independently of protein content, there was no significant difference in the enzymatic activity of either Na^+^/K^+^-ATPase (unpaired t-test; *P* = 0.52; Figure 6B) or V-type H^+^-ATPase (*P* = 0.32) between flies from different acclimation temperatures. In contrast, total protein did not differ between midguts (unpaired t-test; *P* = 0.41) or hindguts (P = 0.30) of warm- and cold-acclimated flies.

**Figure 6.**
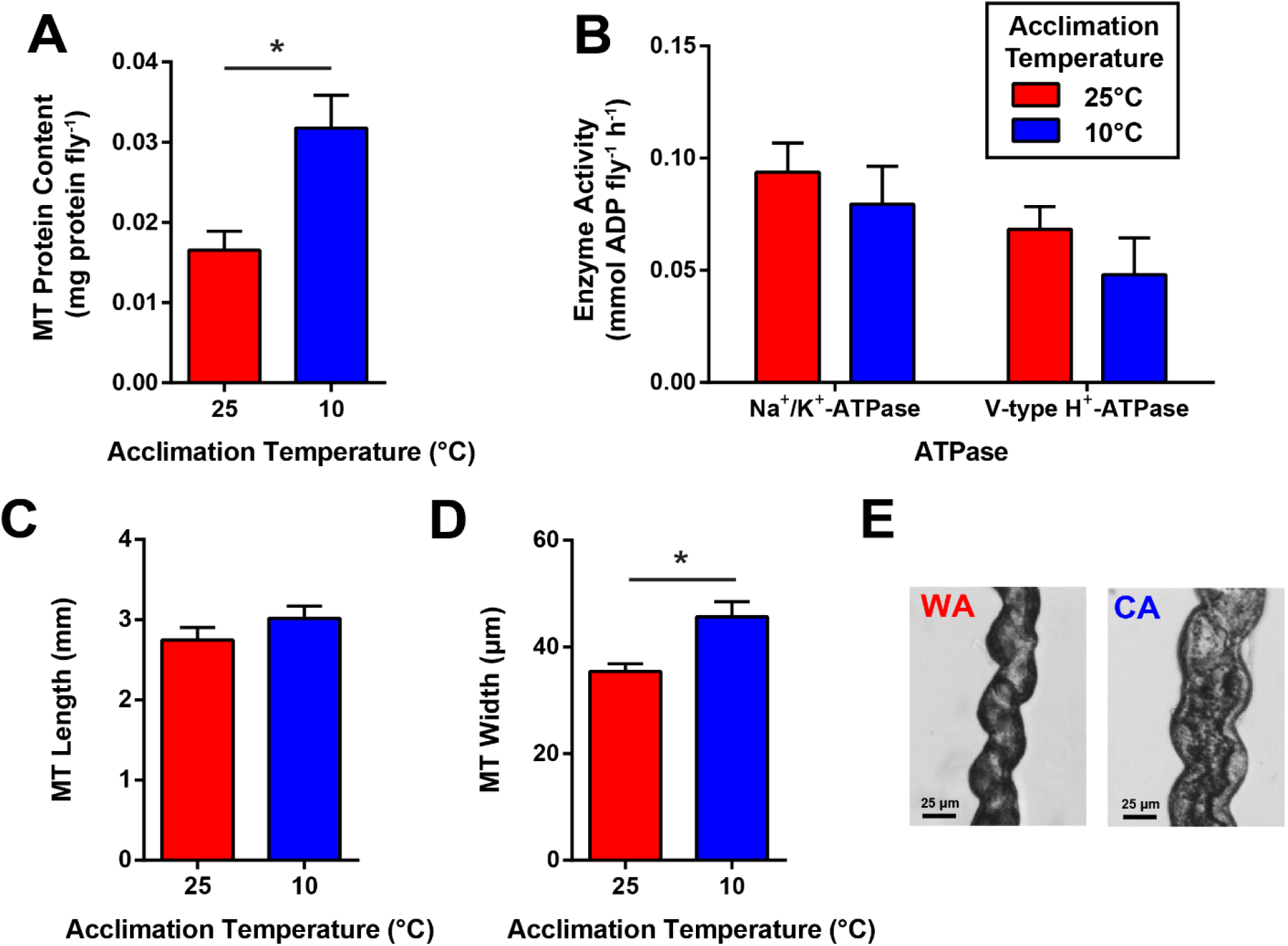
Increased Malpighian tubule width and protein content underlies apparent changes in ion-motive ATPase activity in cold-acclimated *D. melanogaster* females. (A) Malpighian tubule protein content per fly in warm- and cold-acclimated flies (n = 5 sets of Malpighian tubules from 30 flies per acclimation group). (B) Enzymatic activity of Na^+^/K^+^-ATPase and V-type H^+^-ATPase per individual fly (as opposed to protein content) in warm- and cold-acclimated flies. Malpighian tubule (C) length and (D) width assessed in warm- (WA) and cold-acclimated (CA) flies (n = 7 tubules per group). (E) Example image of warm-acclimated (left) and cold-acclimated (right) Malpighian tubule illustrating alterations in MT size. Bars represent mean ± SEM. Asterisk denotes a significant difference (unpaired t-test; *P* < 0.01).

To assess whether the difference in protein content may be driven by difference in Malpighian tubule size, the width and length of Malpighian tubules of flies from both acclimation groups were assessed. Malpighian tubule length did not differ between the two acclimation groups (unpaired t-test; *P* = 0.240; Figure 6C), but the Malpighian tubules of cold-acclimated flies were significantly wider than those of warm-acclimated flies (unpaired t-test; *P* = 0.007; Figure 6D-E).

## 4. Discussion

### 4.1 Cold acclimation mitigates hyperkalemia and improves the cold tolerance of female *D. melanogaster*

Cold acclimation improves the cold tolerance of *D. melanogaster* females and mitigates the degree of cold-induced hyperkalemia. Specifically, cold acclimated flies entered chill coma at a lower temperature (lower CT_min_) and recovered faster from a chill coma (lower CCRT; Figure 2A-B). These results are consistent with recent findings that chill tolerance dramatically improves under the same acclimation treatment, with the Lt_50_ nearly doubling (MacMillan et al., 2017), and where improvements in chill tolerance (CCRT, CT_min_, and survival) have been illustrated in a variety of insects following cold acclimation, including *D. melanogaster* (Andersen et al., 2017a; Koštál et al., 2011; Overgaard et al., 2008; Ransberry et al., 2011). Also consistent with findings in firebugs, crickets, cockroaches, and fruit flies, cold-induced hemolymph [K^+^] elevations were greatly mitigated following cold acclimation (Koštál et al., 2004; Koštál et al., 2006; MacMillan et al., 2015c). This is consistent with the observed improvements in CCRT and survival, that have consistently been related to the prevention of cold-induced hyperkalemia (Koštál et al., 2004; Koštál et al., 2006; MacMillan et al., 2015a; Yerushalmi et al., 2016). Thus, our population of *D. melanogaster* responds to a cold acclimation treatment similarly to previous studies on this species (with improved chill tolerance and mitigated cold-induced hyperkalemia). We therefore assessed whether the ionoregulatory organs of *D. melanogaster* are involved in facilitating this preservation of K^+^ balance.

### 4.2 Physiological plasticity of the Malpighian tubules improves K^+^ clearance

In *Drosophila*, the Malpighian tubules are responsible for the formation of the primary urine and act as the main excretory organ, and thus play a central role in organismal iono- and osmoregulation. We used Ramsay assays and measured secreted fluid ion concentrations with ion-selective microelectrodes to assess temperature effects on fluid, Na^+^, and K^+^ secretion following thermal acclimation. As predicted, the Malpighian tubules of cold-acclimated flies maintained K^+^ secretion rates at low temperatures better than those from warm-acclimated flies, which would facilitate K^+^ clearance in the cold (Figure 3E). This resulted in preserving the [Na^+^]:[K^+^] ratio of the secreted fluid from tubules of cold-acclimated flies, while warm-acclimated flies experienced a 4-fold increase in the [Na^+^]:[K^+^] ratio (Figure 3F). Reduced K^+^ secretion in the tubules of warm-acclimated flies would reduce their capacity for K^+^ clearance and thus likely contribute to the accumulation of hemolymph [K^+^].

The Malpighian tubules are energized by temperature-sensitive ATPases, and the basal rate of fluid secretion across the Malpighian tubules dramatically decreases in the cold (MacMillan et al., 2015d; Ramsay, 1954). In an apparent compensatory response, following one week at 10°C, the fluid secretion rate of tubules from cold-acclimated flies was elevated at 5°C, 10°C, and 25°C, but not at 0°C, relative to warm-acclimated flies (Figure 3A). For instance, the secretion rate of tubules from cold-acclimated flies increased by a factor of ~9 from 0.05 nL min^−1^ to 0.46 nL min^−1^ at their acclamatory temperature of 10°C. These rates are still below those of tubules from warm-acclimated flies at room temperature (~50%) demonstrating that exposure to 10°C for 1-week results in partial compensation of fluid secretion rates. However, at 0°C, the temperature used for CCRT and a previous survival analysis (MacMillan et al., 2017), there was no difference in fluid secretion rates but ion secretion rates differed.

To our knowledge, this is the first assessment of the role of Malpighian tubules in *D. melanogaster* cold acclimation, but both MacMillan *et al*. (2015d) and Andersen *et al*. (Andersen et al., 2017b) assessed the role of the tubules in cold tolerance among five *Drosophila* species raised under common garden conditions (21-22°C). Together, these results suggest that thermal acclimation and adaptation may work through shared or similar physiological mechanisms in the Malpighian tubules. Both cold adaptation and acclimation reduce (or entirely prevent) hemolymph [K^+^] disturbance in the cold (MacMillan et al., 2015d; MacMillan et al., 2017). In both cases more cold tolerant flies better maintained tubule K^+^ secretion and consequently the ratio of [Na^+^]:[K^+^] while warm adapted or acclimated flies experienced preferential secretion of Na^+^ over K^+^ (Andersen et al., 2017c; MacMillan et al., 2015d). Thus, chill tolerance in cold adapted and acclimated flies appears to be at least partially improved by improved tubule K^+^ secretion at low temperature. In addition, in contrast to the tubule fluid secretion in cold acclimated *D. melanogaster*, a recent study on cold-acclimated crickets found reduced fluid secretion rates at lower temperatures, suggesting a different mechanism exists in the cold acclimation of these two insects (Des Marteaux et al., 2018).

### 4.3 Rectal K^+^ reabsorption is lower in cold-acclimated flies

Whereas the Malpighian tubules serve as the primary site for ion and fluid excretion into the gut, the midgut and hindgut absorb ions and water from the gut into the hemolymph (D’Silva et al., 2017; Larsen et al., 2014). As such, we hypothesized that reduced K^+^ absorption in the cold across these epithelia would alleviate cold-induced hyperkalemia. To assess K^+^ absorption across the gut we utilized the scanning ion selective electrode technique to measure K^+^ flux at 6°C and 23°C across the midgut and hindgut (ileum and rectum). While neither acclimation nor exposure temperature impacted mean K^+^ flux in the midgut, differences were observed in the hindgut. Temperature had no effect on the K^+^ flux at the ileum of cold-acclimated flies but ileal K^+^ flux of warm-acclimated flies was higher at 23°C compared to 6°C (Figure 4B). The greatest effect of both acclimation and exposure temperatures was observed in the rectum, where K^+^ flux in the rectums of cold-acclimated flies was significantly lower than that of warm-acclimated flies at both temperatures (Figure 4C). Additionally, K^+^ fluxes at the rectum were reduced at 6°C regardless of acclimation group and this is consistent with reduced metabolic demand in the rectum of grasshoppers in the cold (Palazzo and August, 1997). The K^+^ flux across rectal pads of five *Drosophila* species was also reduced in low temperatures regardless of degree of chill tolerance of the species (Andersen et al., 2017c). The reduction of rectal K^+^ reabsorption at low temperatures likely assists in preventing lethal hyperkalemia, and is thus consistent with the hypothesis of the current study that cold acclimation mitigates hyperkalemia by reducing net K^+^ uptake in the main ionoregulatory epithelia of *D. melanogaster*.

### 4.4 Hemolymph K^+^ balance depends on the integrated functions of the Malpighian tubule and the rectum

Under homeostatic conditions, the Malpighian tubules and rectum, as the primary sites of K^+^ transport, must work in a synchronous and complementary manner to maintain organismal K^+^ balance. Thus, to estimate the degree of cold-induced disruption to whole body K^+^ balance, a comparison between their relative inhibition at low temperature is informative; a mismatch in the effects of temperature on these two organs would lead to an imbalance in hemolymph [K^+^] in the cold. To do so, we first compared the temperature effect on K^+^ transport between 23°C and 6°C for the rectum and between 23°C and 5°C in the Malpighian tubules. This was done by calculating the temperature coefficient (Q_10_), which represents the effect of a 10°C reduction in temperature on K^+^ transport. Q_10_ values were calculated using the following equation (the Malpighian tubules are used as an example):

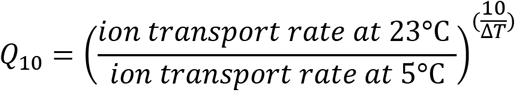

Where ΔT is the difference in temperature (18°C). To estimate the relative effect of temperature on K^+^ transport in the Malpighian tubules in comparison to the rectum, the following equation was used:

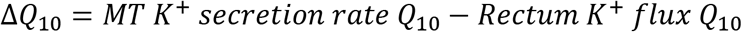

The further this metric deviates from one, the greater imbalance in the overall circuit of K^+^ transport between these two organs. In warm acclimated flies, the Q_10_ of Malpighian tubule K^+^ secretion is 4.7 and the Q_10_ of rectal K^+^ flux is 1.8, resulting in a ΔQ_10_ of 2.7. Hence the cold has a higher effect on Malpighian tubule function than on the rectum. In cold acclimated flies, the Q_10_ of Malpighian tubule K^+^ secretion is 3.2 and the Q_10_ of rectal K^+^ flux is 2.0, resulting in a ΔQ_10_ of 1.2. Therefore, even with cold acclimation, the cold has a greater effect on Malpighian tubule function compared to the rectum; however, cold acclimation reduces the imbalance in K^+^ transport between the two organs such that K^+^-clearance is more severely affected in the cold in comparison to rectal K^+^ absorption in warm acclimated flies compared to cold acclimated flies. As a result, we would expect that K^+^ would accumulate in the hemolymph of warm acclimated flies more rapidly, as observed in the current study. These results are also informative in that they show that the Malpighian tubules are more temperature sensitive than the rectum and, exhibit a greater degree of adjustment to K^+^ transport following cold acclimation both in terms of thermal sensitivity and in absolute K^+^ transport rates. This supports the idea that reduced Malpighian tubule K^+^ clearance is a central problem for *D. melanogaster* at low temperatures, as is the case for crickets (Des Marteaux et al., 2018), and that its preservation is thus beneficial to the development of chill tolerance.

### 4.5 The effects of cold acclimation on ion transport are independent of V-type H^+^- and Na^+^/K^+^-ATPase activity

We hypothesized that cold acclimation would mitigate the loss of ion balance through the alteration of active ion transport in ionoregulatory organs. To test this, we measured the activities of V-type H^+^-ATPase and Na^+^/K^+^-ATPase in isolated midguts, Malpighian tubules, and hindguts of warm- and cold-acclimated flies.

In *D. melanogaster*, an apically-located V-type H^+^-ATPase is the primary driver of Malpighian tubule fluid secretion, while a basolateral Na^+^/K^+^-ATPase contributes 10-19% of fluid secretion and is responsible for increasing the [K^+^]:[Na^+^] ratio in the secreted fluid (Linton and O’Donnell, 1999). Thus, to account for increased Malpighian tubule fluid secretion and the preservation of K^+^ secretion over a variety of temperatures, we expected that the activities of both ion-motive ATPases would increase in cold-acclimated flies. Contrary to this hypothesis, however, the activity of both ATPases significantly decreased in cold-acclimated flies when compared to their warm-acclimated counterparts when these activities were standardized by total protein content of tubules (Figure 5A-B). However, when ATPase activities were expressed per individual organ, no significant differences in activities were found (Figure 6B). Each sample contained the same number of tubules, so we measured Malpighian tubule size and found that cold-acclimated tubules are significantly wider than warm-acclimated tubules (Figure 6D-E). Similar measurements of the Malpighian tubules revealed no such size differences in chill tolerant *Drosophila* species when corrected for total body mass (Andersen et al., 2017b). Currently, it is unclear whether this increased tubule width in cold-acclimated flies stems from hypertrophy, hyperplasia, or simply the enlargement of the Malpighian tubule lumen, and whether this size difference is indeed the cause of increased protein content. Further studies are also required to elucidate whether these morphological changes have any functional relevance. Regardless, we found no evidence to support our hypothesis that functional changes in ion transport would be mediated by changes in ion-motive ATPase activity, suggesting that an alternative mechanism is responsible for the functional differences (e.g. increased fluid secretion and K^+^ transport rates) that were observed in the Malpighian tubules of cold-acclimated flies (discussed below). This is of particular interest, because in crickets, Na^+^/K^+^-ATPase activity increased in the Malpighian tubules despite a decrease in fluid secretion in the cold (Des Marteaux et al., 2018).

At the ureter-gut junction, Malpighian tubule and midgut contents mix prior to their entry into the hindgut, where ions are actively reabsorbed into the hemolymph, resulting in concentrated excreta (Larsen et al., 2014). Unlike the Malpighian tubules, however, our basic mechanistic understanding of insect rectal function is weak. In cockroaches, locusts, and flies, the main site of ion reabsorption occurs at four areas of thickened epithelia known as rectal pads (Larsen et al., 2014). Studies of ion transport across locust rectum suggest that an apical V-type H^+^-ATPase at least partially energizes epithelial ion transport (Gerencser and Zhang, 2003; Phillips et al., 1996). Na^+^/K^+^-ATPase is also highly expressed in *D. melanogaster* hindgut, second in abundance only to the Malpighian tubules, and has been localized to the basolateral membrane in *A. aegypti*, yet its function remains unknown (Chintapalli et al., 2007; Patrick et al., 2006). As is the case in crickets (Des Marteaux et al., 2018), the activities of Na^+^/K^+^- and V-type H^+^-ATPase in the hindgut did not differ between the acclimation treatments in the current study, suggesting an alternative mechanism is responsible for altered K^+^ flux across the ileum and rectum. As with the tubules and hindgut, a mismatch also exists between ion-motive ATPase activity and midgut K^+^ flux. While mean midgut K^+^ flux did not change with acclimation treatment, both Na^+^/K^+^- and V-type H^+^-ATPases decreased in activity.

Cumulatively, these misalignments between ion-motive ATPase activity and ion transport suggest that other mechanisms affect epithelial transport following cold acclimation. Such alternative mechanisms may include: (1) changes to the plasma membrane environment known to affect key transport proteins such as Na^+^/K^+^-ATPase (reviewed by Hazel (1995)), (2) changes in paracellular permeability which may mediate cold-induced ion leak in the cold (Andersen et al., 2017c; MacMillan et al., 2017), (3) changes in endocrine control of ion- and osmoregulation (Terhzaz et al., 2015), (4) changes in mitochondrial ATP production in the cold (Colinet et al., 2017) or (5) changes in the thermal sensitivity of Na^+^/K^+^- or V-type H^+^-ATPase. Since all enzyme activity assays in this study were conducted at 25°C, the thermal sensitivity of these enzymes was not determined here. However, MacMillan (2015b) previously showed that no difference in the thermal sensitivity of Na^+^/K^+^-ATPase exists following cold-acclimation in *D. melanogaster*. The thermal sensitivity of V-type H^+^-ATPase following cold acclimation remains a possibility that should be assessed in future studies.

#### Conclusion

As previously demonstrated in a variety of chill-susceptible insects, cold acclimation led to reduced CT_min_, faster recovery from a chill coma, and reduced degree of cold-induced hemolymph [K^+^] in *D. melanogaster*. This improvement in hemolymph K^+^ balance in the cold coincided with increased Malpighian tubule K^+^ and fluid secretion rates at low temperatures. In parallel, reabsorption of K^+^ was reduced in the rectum but unchanged in the midgut of cold-acclimated flies in comparison to warm-acclimated flies. Together, these changes illustrate that cold-acclimated flies have a greater capacity for K^+^ clearance than warm-acclimated flies in the cold and support an important role for these ionoregulatory organs in the prevention of cold-induced hyperkalemia following cold acclimation. Furthermore, measurement of the activities of Na^+^/K^+^-ATPase and V-type H^+^-ATPase revealed no clear link to K^+^ transport across the midgut, Malpighian tubules, or hindgut, suggesting that modulation of these organs following cold acclimation is mediated through an alternative mechanism. Our results lend support to the role of plasticity of the Malpighian tubule and the rectum in the cold acclimation of chill-susceptible insects, and the independence of this functional plasticity to the modulation of Na^+^/K^+^-ATPase and V-type H^+^-ATPase.

## Acknowledgments

The authors would like to thank Sima Jonusaite, Andrea Durant and Fargol Nowghani for training and advice on enzyme activity assays. The authors also thank Dr. Scott Kelly and Dr. Carol Bucking for providing access to required laboratory instruments and Dr. Paluzzi for his feedback throughout the project.

## Competing interests

The authors declare no competing interests.

## Author contributions

GY, HAM, and AD conceived of the study. GY, LM, and HAM collected the data. GY analyzed the data and drafted the manuscript, and all authors edited the manuscript.

## Funding

This work was supported by a Natural Sciences and Engineering Research Council of Canada (NSERC) Discovery grant to AD, an NSERC Banting Postdoctoral Fellowship to HAM, an NSERC Alexander Graham Bell Canada Graduate Scholarship (CGS-M) to GY and York University Faculty of Science Stong Scholarships to GY and LM.

## Figures

**Table S1.** Results of two-way ANOVAs assessing the effects of acclimation and exposure temperatures on Malpighian tubule function. Ramsay assays and ion-selective microelectrodes were used to measure fluid secretion rate, [Na^+^] in secreted fluid, [K^+^] in secreted fluid, Na^+^ secretion rate, K^+^ secretion rate, and the ratio of Na^+^:K^+^ in the secreted fluid. DF = degrees of freedom.

| **Trait** | **Variable** | **Statistic (DF)** | **P-value** |
| --- | --- | --- | --- |
| Secretion Rate | Acclimation temperature x Exposure temperature | F (3, 70) = 6.6 | P = 0.0006 |
|  | Exposure temperature | F (3, 70) = 91.6 | P < 0.0001 |
|  | Acclimation temperature | F (1, 70) = 21.5 | P < 0.0001 |
| [Na<sup>+</sup>] in Secreted Fluid | Acclimation temperature x Exposure temperature | F (3, 66) = 5.7 | P = 0.0016 |
|  | Exposure temperature | F (3, 66) = 22.5 | P < 0.0001 |
|  | Acclimation temperature | F (1, 66) = 54.2 | P < 0.0001 |
| [K<sup>+</sup>] in Secreted Fluid | Acclimation temperature x Exposure temperature | F (3, 67) = 3.7 | P = 0.0157 |
|  | Exposure temperature | F (3, 67) = 7.6 | P = 0.0002 |
|  | Acclimation temperature | F (1, 67) = 17.3 | P < 0.0001 |
| Na<sup>+</sup> Secretion Rate | Acclimation temperature x Exposure temperature | F (3, 68) = 4.5 | P = 0.0065 |
|  | Exposure temperature | F (3, 68) = 117.1 | P < 0.0001 |
|  | Acclimation temperature | F (1, 68) = 5.1 | P = 0.0265 |
| K<sup>+</sup> Secretion Rate | Acclimation temperature x Exposure temperature | F (3, 69) = 3.3 | P = 0.0239 |
|  | Exposure temperature | F (3, 69) = 57.0 | P < 0.0001 |
|  | Acclimation temperature | F (1, 69) = 12.1 | P = 0.0009 |
| Na<sup>+</sup>:K<sup>+</sup> Ratio in Secreted Fluid | Acclimation temperature x Exposure temperature | F (3, 66) = 6.8 | P = 0.0005 |
|  | Exposure temperature | F (3, 66) = 7.7 | P = 0.0002 |
|  | Acclimation temperature | F (1, 66) = 28.3 | P < 0.0001 |

